# GpsB control of PASTA kinase activity in *Listeria monocytogenes* influences peptidoglycan synthesis during cell wall stress and cytosolic survival

**DOI:** 10.1101/2023.06.12.544644

**Authors:** Jessica L. Kelliher, McKenzie E. Daanen, John-Demian Sauer

## Abstract

The ability to respond quickly to changing environmental conditions in the host is critical for bacterial pathogens. Penicillin-binding protein and serine/threonine-associated (PASTA) kinases are a conserved family of kinases important for cell envelope stress responses in *Firmicutes* and *Actinobacteria*, including the cytosolic pathogen *Listeria monocytogenes*. As serine/threonine kinases, PASTA kinases phosphorylate multiple substrates, yet the mechanisms through which these substrates promote resistance to cell wall stress remain poorly understood. We previously identified GpsB as a target of PrkA, the PASTA kinase in *L. monocytogenes*, through a phosphoproteomics screen. Here, we demonstrate that GpsB can be directly phosphorylated by PrkA, and that mutation of the PrkA-dependent phosphosite T88 to a phosphoablative residue enhances PrkA activity *in vitro*. We find that relative to a strain of *L. monocytogenes* harboring the *gpsB*_T88A_ allele, a strain with the phosphomimetic *gpsB*_T88D_ allele is more sensitive to the cephalosporin antibiotic ceftriaxone and has a diminished capacity to increase peptidoglycan synthesis during stress. We find that GpsB-dependent control of PrkA activity is required for optimal survival and replication of *L. monocytogenes* in the macrophage cytosol. Finally, we show that GpsB is required for full virulence of *L. monocytogenes*, due in part to its role in modulating PrkA activity. Cumulatively, these results demonstrate that phosphorylative feedback between GpsB and PrkA is important for the ability of *L. monocytogenes* to respond to cell wall stress, survive in its cytosolic niche, and cause infection.

## INTRODUCTION

Rising rates of antibiotic resistance among bacterial pathogens is an increasingly urgent threat to human health (1). Many currently used antibiotics target bacterial cell wall synthesis, but bacteria can be resistant to such drugs through acquired or intrinsic mechanisms. While much effort has been focused on understanding acquired mechanisms of resistance, knowledge of intrinsic mechanisms, such as cell wall stress response systems, is also critical to combatting the threat of antimicrobial resistance. An important intrinsic resistance determinant conserved in *Firmicutes* and *Actinobacteria* is the penicillin-binding protein and serine/threonine-associated, or PASTA, kinase family. PASTA kinases have extracellular PASTA domains that sense perturbations in the cell envelope and mediate a global stress response via their cytoplasmic eukaryotic-like serine/threonine kinase domain, phosphorylating multiple downstream targets involved in cell wall homeostasis, central metabolism, and more (2–6). In a variety of bacteria, including important pathogens like *Listeria monocytogenes*, *Staphylococcus aureus*, *Mycobacterium tuberculosis*, *Enterococcus faecalis*, and *Streptococcus pneumoniae*, PASTA kinases are important for growth in the presence of cell wall-targeting agents (e.g., β-lactams and lysozyme) and also critically contribute to pathogenesis (7–14). The use of PASTA kinase-specific inhibitors has been explored as a potential antibiotic adjuvant, including in organisms with additional acquired resistance mechanisms (15–18).

*Listeria monocytogenes* is a ubiquitous Gram-positive saprophyte that can be pathogenic in mammals. The systemic disease listeriosis is of particular concern for young, old, pregnant, or immunocompromised individuals and has a high mortality rate (20-30%) despite currently effective antibiotic treatment options (19, 20). The well-defined pathogenic lifecycle of *L. monocytogenes* involves entry into cells via either internalin-mediated invasion or phagocytosis, escape from the vacuole using the pore-forming cytolysin LLO, survival and replication in the cytosol, and cell-to-cell spread using actin motility (21). To survive in the cytosol, pathogens like *L. monocytogenes* must resist cell autonomous defense mechanisms that restrict the growth of non-adapted bacteria (22, 23). Although the cytosol is known to be antimicrobial, the molecular details underpinning restriction of bacterial growth and in turn the adaptations used by cytosolic pathogens to survive in this niche are not well understood. Continued investigation of those host-pathogen interface using model pathogens such as *L. monocytogenes* has the potential to reveal avenues for both host- and pathogen-directed therapeutic interventions.

We have previously shown that the PASTA kinase in *L. monocytogenes*, PrkA, is required for survival in the macrophage cytosol (7, 24). A phosphoproteomics screen revealed that during cell wall stress, PrkA phosphorylates 23 proteins involved in diverse physiologic processes, including cell wall synthesis, central metabolism, oxidative stress, and more (24). Targeted genetic analysis revealed that PrkA phosphorylation of ReoM, a small protein that modulates stability of the peptidoglycan (PG) synthesis enzyme MurA (25), drives an increase in PG synthesis in the context of cell wall stress (24). We determined that PrkA-mediated regulation ReoM is required for growth in the presence of cell wall stress, cytosolic survival, and virulence of *L. monocytogenes*. Yet, how the other ∼20 proteins phosphorylated by PrkA contribute to the intrinsic response to cell wall stress and to adaptation to the cytosol remains unknown.

Another putative PrkA substrate identified via phosphoproteomics with a function in cell wall homeostasis is GpsB, a small protein conserved in the *Firmicutes* phylum (26, 27). GpsB was first described in *Bacillus subtilis* for its role in <u>g</u>uiding <u>p</u>enicillin-binding protein (PBP) <u>s</u>huttling between the divisome at the septum and the elongasome at the lateral side wall according to the bacterial cell cycle (26). GpsB also has been shown to interact with multiple proteins involved in cell division and cell wall metabolism in several species, including *B. subtilis*, *L. monocytogenes*, and *S. pneumoniae* (27–30). A *L. monocytogenes* Δ*gpsB* mutant has a growth defect at elevated temperatures and is sensitive to the peptidoglycan (PG)-targeting insults penicillin and mutanolysin (27). These phenotypes can be rescued by deletion of PBP A1, with which GpsB is known to interact, suggesting that GpsB has a role in regulating PBP A1 activity in *L. monocytogenes* (27, 31). Interestingly, in *B. subtilis*, *S. pneumoniae*, and *Enterococcus faecalis*, GpsB homologs have a second described function: they interact with the PASTA kinase in those organisms to modulate kinase activity (32–34). In *B. subtilis* and *E. faecalis*, GpsB is itself a substrate of the PASTA kinase; non-phosphorylated GpsB potentiates kinase activity while the phosphorylated form diminishes it, suggesting that GpsB and the PASTA kinase form a negative feedback loop (32, 34). A previous report has suggested that the phosphorylation state of residue T88 of *L. monocytogenes* GpsB, which we identified as a putative PrkA-dependent phosphosite (24), affects growth at high temperature, with a phosphoablative T88A but not phosphomimetic T88D version rescuing growth of a Δ*gpsB* mutant (35). However, the role of GpsB in modulating activity of the PASTA kinase in *L. monocytogenes* has not been explored.

Here, we demonstrate that GpsB is a direct target of PrkA and can modulate kinase activity *in vitro*, with a T88A version of GpsB enhancing PrkA activity more effectively than wild-type GpsB. We find that in *L. monocytogenes*, a phosphomimetic GpsB_T88D_ dampens growth and peptidoglycan synthesis in the presence of cell wall-targeting antibiotics, indicating that proper phosphoregulation of GpsB is critical during cell wall stress. We find that the ability of GpsB to modulate PrkA activity is important for the ability of *L. monocytogenes* to survive and replicate in the macrophage cytosol. Finally, we show that GpsB is required for the virulence of *L. monocytogenes*, and that this requirement is partially attributable to its role in modulating PrkA activity. Cumulatively, our results demonstrate that the primary function of GpsB in *L*. *monocytogenes* is regulation of the PASTA kinase PrkA, and that this regulation is important for the ability of *L. monocytogenes* to adapt to its infectious niche.

## RESULTS

### GpsB is a substrate of PrkA and potentiates its activity

GpsB was identified as a putative phosphosubstrate of PrkA in *L. monocytogenes* by phosphoproteomics (24) and has been confirmed as a target of the PASTA kinases in *Bacillus subtilis* (32) and *Enterococcus faecalis* (6). To validate that GpsB is a direct target of PrkA in *L. monocytogenes*, we purified GpsB and the intracellular kinase domain and juxtamembrane linker region of PrkA (residues 1-338) and assessed PrkA-dependent GpsB phosphorylation in an *in vitro* kinase assay. While PrkA autophosphorylation and PrkA-dependent phosphorylation of the generic serine/threonine substrate MBP was observed in an endpoint assay, GpsB phosphorylation was not detected in these conditions (Fig. 1A). Previous work in *B. subtilis*, *S. pneumoniae*, and *E. faecalis* has demonstrated that the presence of GpsB can potentiate PASTA kinase-dependent phosphorylation of other substrates (32–34). Therefore, we assessed how addition of GpsB to a kinase reaction with MBP would affect phosphorylation of MBP. In an end-point assay, MBP was phosphorylated to a similar extent regardless of whether GpsB was also in the reaction (Fig. 1B). In these reaction conditions, phosphorylated GpsB was also detected, indicating that GpsB can be directly phosphorylated by PrkA.

**Figure 1.**
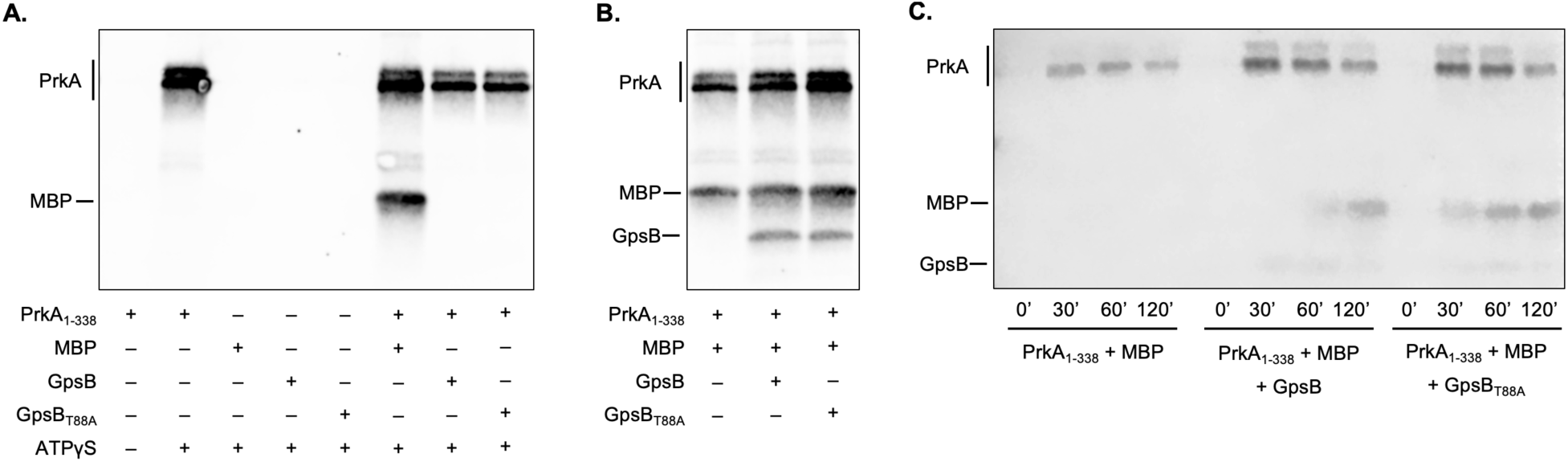
GpsB is a substrate of PrkA and potentiates its activity. (**A**-**B**) Endpoint *in vitro* kinase assays of recombinantly purified PrkA_1-338_ (kinase domain plus juxtamembrane linker) activity against the generic serine/threonine kinase substrate MBP and/or recombinantly purified GpsB or GpsB_T88A_. The indicated combinations of proteins, MBP, and ATP analog ATPγS were incubated overnight at room temperature, followed by a reaction to alkylate thiophosphorylated sites and Western blotting with an anti-thiophosphate ester antibody. ATPγS was present in all reactions in panels **B** and **C**. (**C**) Time-course *in vitro* kinase assays of PrkA_1-338_ activity against the MBP in the absence and presence of recombinantly GpsB or GpsB_T88A_. The indicated combinations of proteins and MBP were incubated at room temperature. At the indicate time points, kinase reactions were quenched with EDTA, followed by alkylation and Western blotting. (**A**-**C**) Blots are representative of at least two biological replicates.

To investigate whether GpsB affects the kinetics of PrkA-dependent phosphorylation of MBP, we next performed a time-course assessment of the reaction. While in the absence of GpsB there was PrkA autophosphorylation but no detectable phospho-MBP within 2 hours, in the presence of GpsB we could detect phospho-MBP within 30 minutes (Fig. 1C). Phosphorylated GpsB was also detected at low levels in this reaction (Fig. 1C).

In *B. subtilis* and *E. faecalis*, the non-phosphorylated form of GpsB increases activity of its cognate PASTA kinase more than the phosphorylated form of GpsB (32, 34). To test whether the phosphorylation status of GpsB affects its ability to regulate PrkA activity in *L. monocytogenes*, we purified a T88A mutant of GpsB and assessed PrkA-dependent MBP phosphorylation in endpoint and kinetic assays as above. Of note, while our previous phosphoproteomics experiment could not definitively call the PrkA-dependent phosphosite from among four residues on GpsB (T71, T73, T88, or T89) (24), T88 corresponds to the PASTA kinase-phosphorylated T75 of *B. subtilis* GpsB (32), and Cleverly *et al.* have previously shown that the phosphorylation status of residue T88 of *L. monocytogenes* GpsB is important for temperature-dependent growth (35). In an endpoint assay with GpsB_T88A_, MBP was phosphorylated to a similar degree as in the reaction with wild-type GpsB (Fig. 1B). However, in a time-course assay, MBP was phosphorylated more robustly in the presence GpsB_T88A_ than in the presence wild-type GpsB (Fig. 1C). Therefore, similar to in *B. subtilis* and *E. faecalis*, a phosphoablative T88A version of GpsB potentiates activity of the PASTA kinase in *L. monocytogenes* more effectively than wild-type GpsB. Cumulatively, these results suggest that the PrkA-dependent phosphorylation status of residue T88 of GpsB can regulate activity of PrkA.

### The phosphoablative GpsB_T88A_ variant promotes growth of *L. monocytogenes* at high temperature

We next took a targeted genetic approach to understand the impact of PrkA-dependent phosphorylation of GpsB on the physiology of *L. monocytogenes*. Previous work has shown that *L. monocytogenes* strain EGD-e mutants lacking *gpsB* grow like wild type at 30°C but have a slower growth rate at 37°C and are unable to grow at 42°C (27). We generated a Δ*gpsB* mutant in the *L. monocytogenes* strain 10403s background and found that the mutant grows similar to wild type at 30°C (Fig. 2A). At 37°C, growth of the Δ*gpsB* mutant was also indistinguishable from that of wild type (Fig. 2B). At 42°C, the Δ*gpsB* mutant had a severe growth defect but was able to complete several doublings, albeit with a dramatically reduced growth rate (Fig. 2C). This observation is in contrast to the findings of Rismondo *et al.* who observed that a Δ*gpsB* mutant is unable to replicate at 42°C (27). The growth defect at 42°C could be complemented by reintroduction of a *gpsB* allele *in trans* under the control of a constitutive, highly expressed promoter (Fig. 2F).

**Figure 2.**
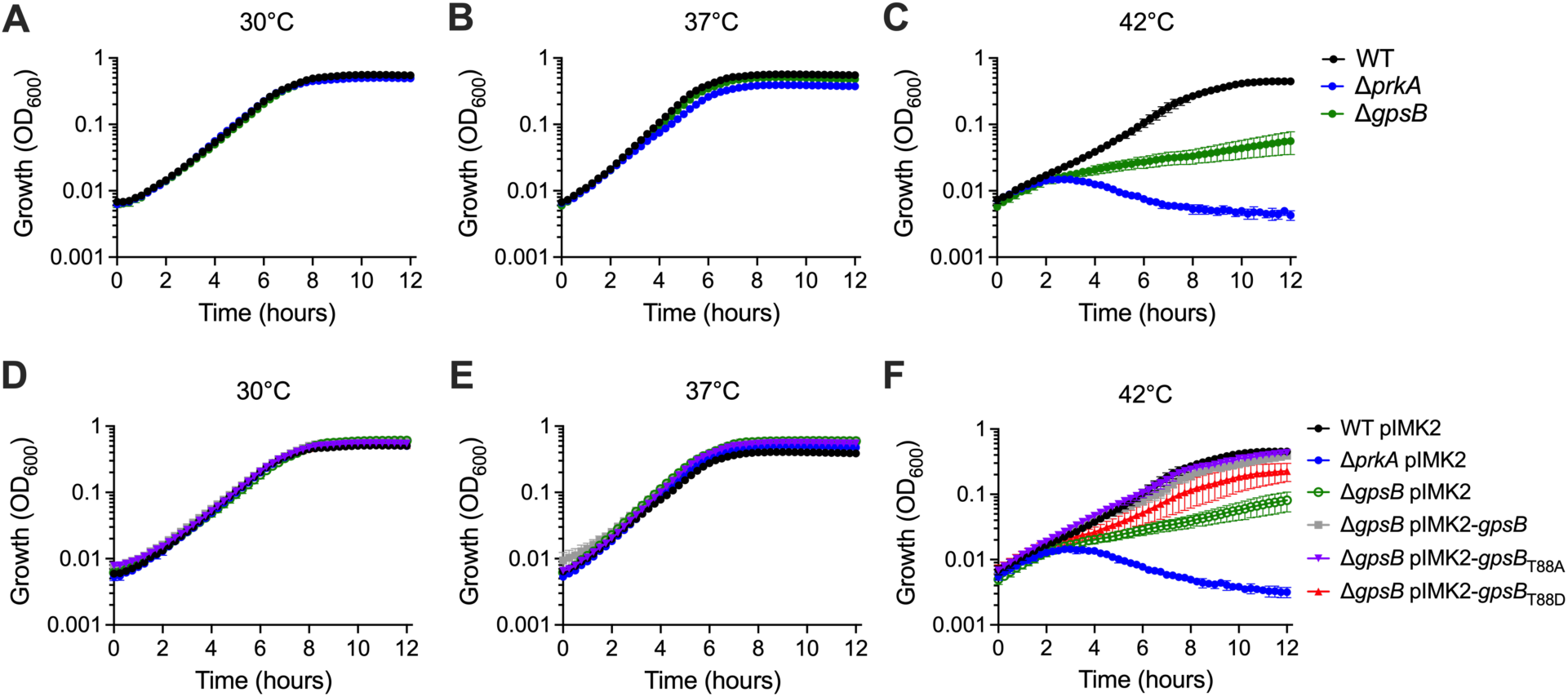
GpsB_T88A_ restores growth of a Δ*gpsB* mutant of *L. monocytogenes* at high temperature. Growth of the indicated strains at 30°C (**A** and **D**), 37°C (**B** and **E**), or 42°C (**C** and **F**) in BHI was monitored by OD_600_. Error bars indicate SEM; n = 3.

While our data demonstrated that a Δ*gpsB* mutant in 10403s was able to grow slowly at 42°C, we observed that a Δ*prkA*::*erm* mutant was unable to grow at this temperature (Fig. S1C, Fig. 2C), similar to the EGD-e Δ*gpsB* mutant (27). Therefore, we next tested the hypothesis that PrkA-dependent phosphorylation of GpsB affects growth of *L. monocytogenes* 10403s at high temperature by reintroducing a phosphoablative (*gpsB*_T88A_) or phosphomimetic (*gpsB*_T88D_) allele *in trans* under the same highly expressed promoter as the wild-type allele. Both mutants grew similarly to wild type at 30°C and 37°C (Fig. 2D-E). However, while the phosphoablative T88A form of GpsB fully restored growth at high temperature, the T88D variant only partially rescued the Δ*gpsB* mutant at 42°C (Fig. 2F). This finding is consistent with prior results by Cleverley *et al.* that shows a T88A but not T88D allele of *gpsB* could fully restore growth of an EGD-e Δ*gpsB* mutant at 42°C (35). Cumulatively, these data suggest that the ability of GpsB to potentiate PrkA activity is important for growth of *L. monocytogenes* at elevated temperatures.

### Phosphorylation status of GpsB affects intrinsic resistance of *L. monocytogenes* to cell wall-targeting antibiotics

Given the role of GpsB in cell division and cell wall homeostasis in *L. monocytogenes* and other bacteria (26–28), we next tested the hypothesis that GpsB is important for intrinsic resistance to cell envelope targeting stressors. In an MIC assay, we found that the Δ*gpsB* mutant is more sensitive than wild type, but not as sensitive as the Δ*prkA* mutant, to the β-lactams ceftriaxone (CRO) and ampicillin, as well as the host-derived, PG-degrading enzyme lysozyme (Table 1, Fig. 3A). The Δ*gpsB* mutant was not more sensitive than wild-type or the Δ*prkA* mutant to the protein synthesis inhibitor chloramphenicol (Table 1), suggesting the role of GpsB is specific to resisting cell wall-targeting antibiotics. The sensitivity of the Δ*gpsB* mutant could be complemented by reintroducing a wild-type copy of *gpsB in trans* (Table 1, Fig. 3A).

**Figure 3.**
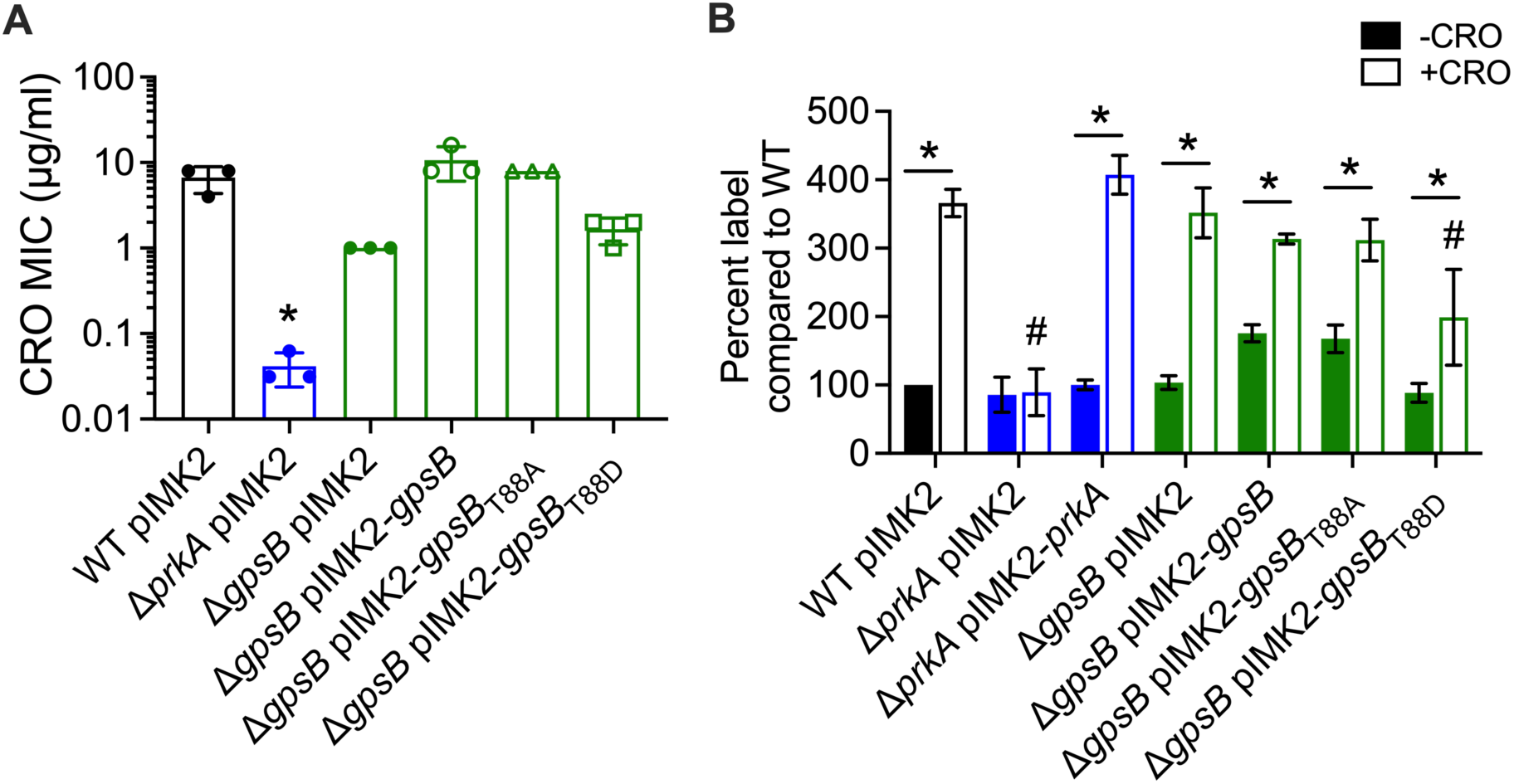
GpsB potentiation of PrkA activity is required for the intrinsic response to cell wall stress. (**A**) Bars indicate median MICs of CRO for the indicated *L. monocytogenes* strains; n = 3. *, *P* < 0.05 compared to wild type by one-way ANOVA with Tukey’s multiple comparisons test. (**B**) The indicated strains of *L. monocytogenes* were grown in the presence of 1 mM EDA-DA for 1 hour in the absence and presence of one-half their respective MIC of CRO. Cells were subjected to a copper-catalyzed alkyne-azide cycloaddition reaction with a modified Alexa Fluor 488, and fluorescence was subsequently analyzed by flow cytometry, with a minimum of 10^5^ events read per sample. Median fluorescence intensities were normalized to that of WT minus CRO. *, *P* < 0.05 for the indicated comparisons by two-way ANOVA with Sidak’s multiple comparisons test. #, *P* < 0.05 compared to wild type plus CRO by two-way ANOVA with Tukey’s multiple comparisons test.

**Table 1.**
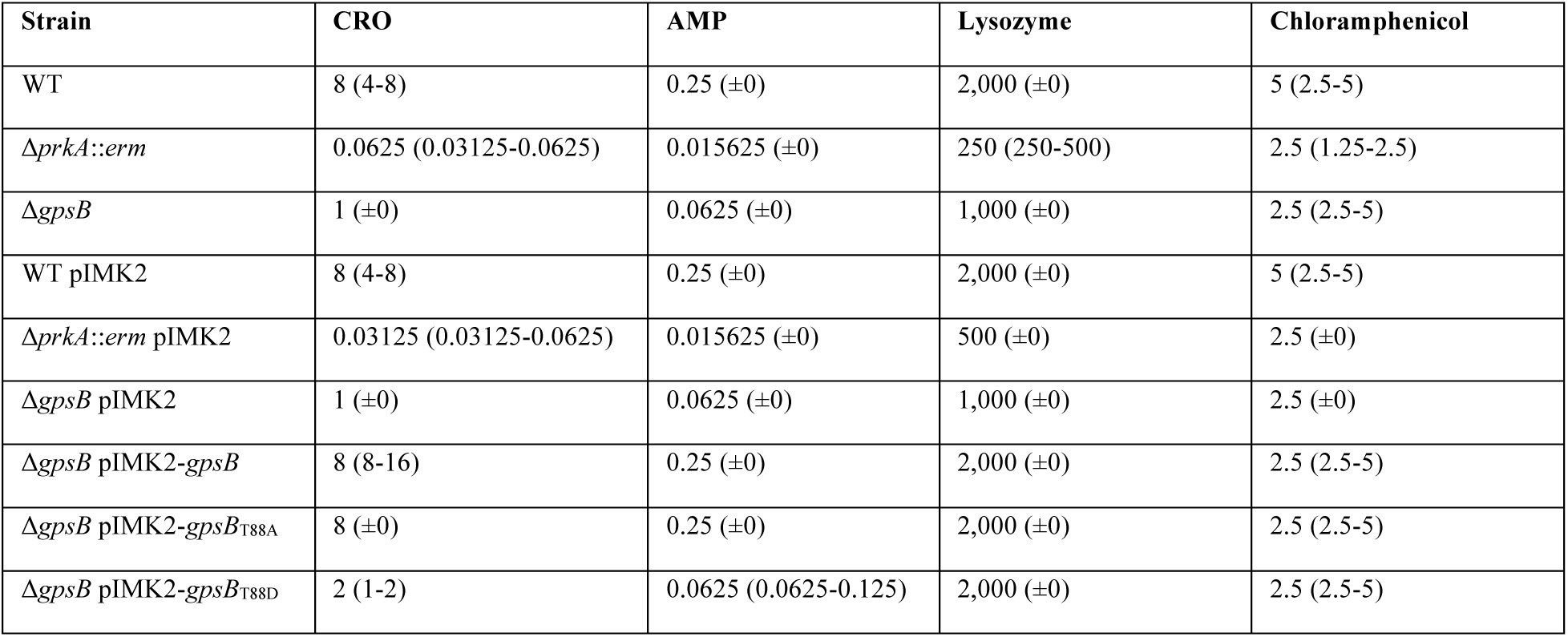
MICs of the indicated compounds against Δ*gpsB* strains in BHI. Values are the median MICs of three biological replicates (and range of values) reporter in μg/ml.

To determine whether the importance of GpsB during cell wall stress is related to its role in control of PrkA activity, we next assessed the ability of a phosphoablative T88A or phosphomimetic T88D allele of *gpsB* to rescue the Δ*gpsB* strain. In the case of CRO, ampicillin, and lysozyme, the T88A allele restored growth to wild-type levels (Fig. 3A, Table 1). In contrast, the T88D allele only partially restored growth of the Δ*gpsB* mutant in the presence of CRO and ampicillin (Fig. 3A, Table 1). Taken together, these observations suggest that the role of GpsB in modulating PrkA kinase activity is necessary for intrinsic resistance to cell wall stressors such as β-lactams.

### A phosphomimetic GpsB allele dampens PrkA-dependent cell wall synthesis during stress

We previously determined that PrkA activity is required for an increase in PG synthesis in *L. monocytogenes* during cell wall stress (24). To investigate how PrkA-dependent phosphorylation of GpsB affects this process, we used an assay described previously to measure PG synthesis in the absence or presence of CRO. In this assay, strains of interest are grown in the presence of an alkyne-conjugated D-alanine-D-alanine analog (EDA-DA) that can be imported, incorporated into nascent PG, and subsequently labeled using a click-chemistry reaction (24, 36). Consistent with prior results, we found that WT and a Δ*prkA* mutant label similarly in the absence of stress, but while WT label significantly increases (∼3-fold) in the presence of CRO, the Δ*prkA* mutant does not exhibit the same increase in label (Fig. 3B), indicating that PrkA is necessary for the increase in PG synthesis during stress. The defect in PG synthesis in the Δ*prkA* mutant in the presence of CRO could be complemented by introduction of *prkA in trans* (Fig. 3B). We observed that the Δ*gpsB* mutant has similar basal levels of labeling as WT in the absence of stress and that labeling also increased in the presence of cell wall stress to a similar magnitude as WT (Fig. 3B). Of note, this finding suggests that interaction with GpsB is not required for the PrkA-dependent increase in PG synthesis during cell wall stress.

We next evaluated the impact of PrkA phosphoregulation of GpsB on PG synthesis in the absence and presence of cell wall stress. The strain constitutively expressing the T88A variant of GpsB exhibited approximately twice as much PG synthesis in the absence of CRO compared to wild-type *L. monocytogenes*, consistent with a model whereby GpsB_T88A_ potentiates PrkA activity even in the absence of cell wall stress (Fig. 3B). Upon the addition of stress, PG labeling in the T88A strain increased to levels similar to wild type plus CRO. In contrast, the strain constitutively expressing the T88D variant of GpsB had similar amounts of PG label as WT in the absence of stress but exhibited significantly lower levels of PG labeling as WT (∼65%) when CRO was added (Fig. 3B), consistent with a model whereby GpsB_T88D_ dampens PrkA activity, preventing optimal PG synthesis during stress. Interestingly, when the Δ*gpsB* mutant was complemented with a constitutively expressed wild-type copy of *gpsB*, PG labeling was ∼2-fold elevated in the absence of CRO (Fig. 3B), suggesting that in the absence of stress, an excess of non-phosphorylated GpsB may lead to aberrant stimulation of PrkA activity. Altogether, these results suggest that while GpsB is not required for PrkA activation, its presence and phosphorylation state can both positively and negatively regulate PrkA-dependent PG synthesis control *L. monocytogenes*.

### GpsB potentiation of PrkA activity is important for survival and replication in the macrophage cytosol

To investigate the importance of GpsB to adaptation of *L. monocytogenes* to the macrophage cytosol, we measured cytosolic killing using a previously described bacteriolysis reporter (37). In this system, *L. monocytogenes* strains carry a plasmid with a CMV promoter-driven luciferase gene that is only expressed when a bacterial cell lyses in the cytosol and the plasmid is translocated to the nucleus. Consistent with prior results, a Δ*prkA* mutant is killed ∼5-10-fold more frequently than wild-type (Fig. 4A). The Δ*gpsB* mutant displayed an intermediate phenotype, lysing substantially more than WT but not as much as Δ*prkA* (Fig. 4A), indicating that GpsB is important for survival in the macrophage cytosol. This defect could be complemented by reintroduction of a wild-type allele of *gpsB* on a plasmid (Fig. 4A). We next evaluated how the phosphorylation status of GpsB impacts cytosolic survival of *L. monocytogenes*. The phosphoablative T88A allele of *gpsB* was able to restore cytosolic survival to wild-type levels (Fig. 4A). In contrast, the phosphomimetic T88D variant of *gpsB* was killed at levels similar to the *gpsB* mutant (Fig. 4A). Overall, these data suggest that the role of GpsB in stimulating PrkA activity is critical for survival in the macrophage cytosol.

**Figure 4.**
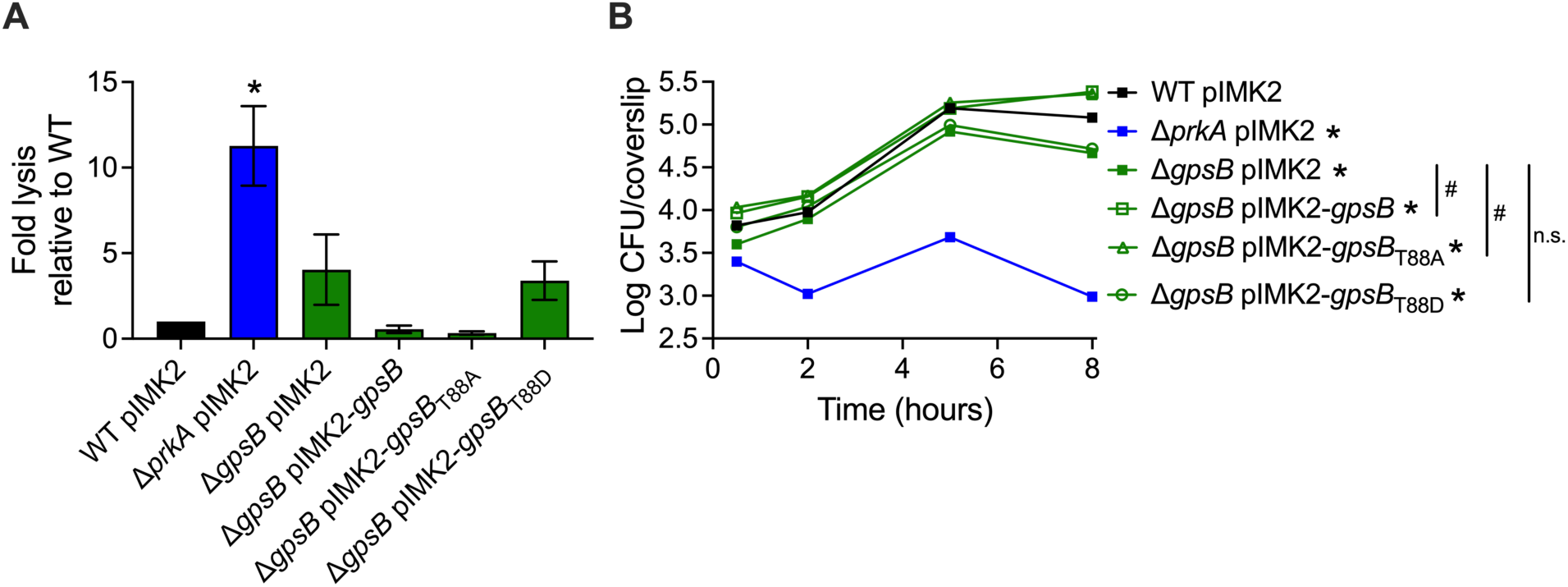
GpsB control of PrkA activity is required for survival and replication in macrophages. (**A**) Intracellular bacteriolysis in immortalized *Ifnar*^-/-^ macrophages. Macrophages were infected with the indicated strains carrying the pBHE573 reporter vector at an MOI of 10, and luciferase activity was measured 6 hours post infection. Error bars indicate SEM; n = 3. (**B**) Intracellular growth curve in BMDMs. BMDMs seeded on coverslips were infected with the indicated strains at an MOI of 0.2. CFUs were enumerated per coverslip in triplicate at the indicated time points. Error bars indicate SD; growth curve is representative of three biological replicates. *, *P* < 0.05 compared to wild type and #, *P* < 0.05 compared to Δ*gpsB* pIMK2 at the 8-hour time point by two-way ANOVA with Tukey’s multiple comparisons test.

Given the importance of the phosphorylation status of GpsB for intracellular survival, we next tested the hypothesis that PrkA-dependent phosphorylation of GpsB is important for the ability of *L. monocytogenes* to replicate in the macrophage cytosol. Consistent with our previous findings (24), while wild-type *L. monocytogenes* replicates robustly in bone marrow-derived macrophages, the Δ*prkA* mutant is unable to grow (Fig. 4B). Although the Δ*gpsB* mutant was able to replicate intracellularly, its terminal burden after 8 hours of growth was slightly but significantly lower than wild type (Fig. 4B). Complementation with either a wild-type copy of *gpsB* and the phosphoablative T88A allele resulted in a growth yield approximately 1 log greater than that of the Δ*gpsB* mutant (Fig. 4B). In contrast, the phosphomimetic T88D strain reached a reached a final burden below wild type, similar to the Δ*gpsB* mutant (Fig. 4B). Altogether, these data suggest that, while GpsB is not absolutely required for intracellular growth, its primary role in the macrophage cytosol is to modulate PrkA activity for optimal cytosolic survival and replication.

### GpsB is required during systemic listeriosis due to PrkA-dependent and -independent roles

We next investigated whether GpsB is important for systemic disease caused by *L. monocytogenes* in a mouse model of listeriosis. Mice were infected intravenously with strains of interest, and bacterial burdens in the liver and spleen were enumerated after two days of infection. As we have observed previously, no bacteria were recovered from either the livers or the spleens of mice infected with the Δ*prkA* mutant (Fig. 5A-B). This virulence defect could be complemented by expression of *prkA* from a constitutive promoter *in trans* (Fig. 5). The Δ*gpsB* mutant was attenuated ∼100-fold in both the liver and the spleen, indicating that GpsB is required for *in vivo* virulence of *L. monocytogenes* (Fig. 5). Because the infections lasted 48 hours at an elevated temperature (∼37°C), we tested the hypothesis that the virulence of the Δ*gpsB* mutant could be due to suppressor mutations arising, since the infection proceeds for 48 hours. We screened 1 colony from the liver and spleen of each mouse infected with the Δ*gpsB* mutant for the ability to grow at 42°C. We found that all isolates were all still sensitive to the elevated temperature (data not shown), suggesting they do not have suppressing mutations in their genomes. Reintroduction of a wild-type allele of *gpsB in trans* partially restored the virulence defect of Δ*gpsB* mutant in both organs (Fig. 5).

**Figure 5.**
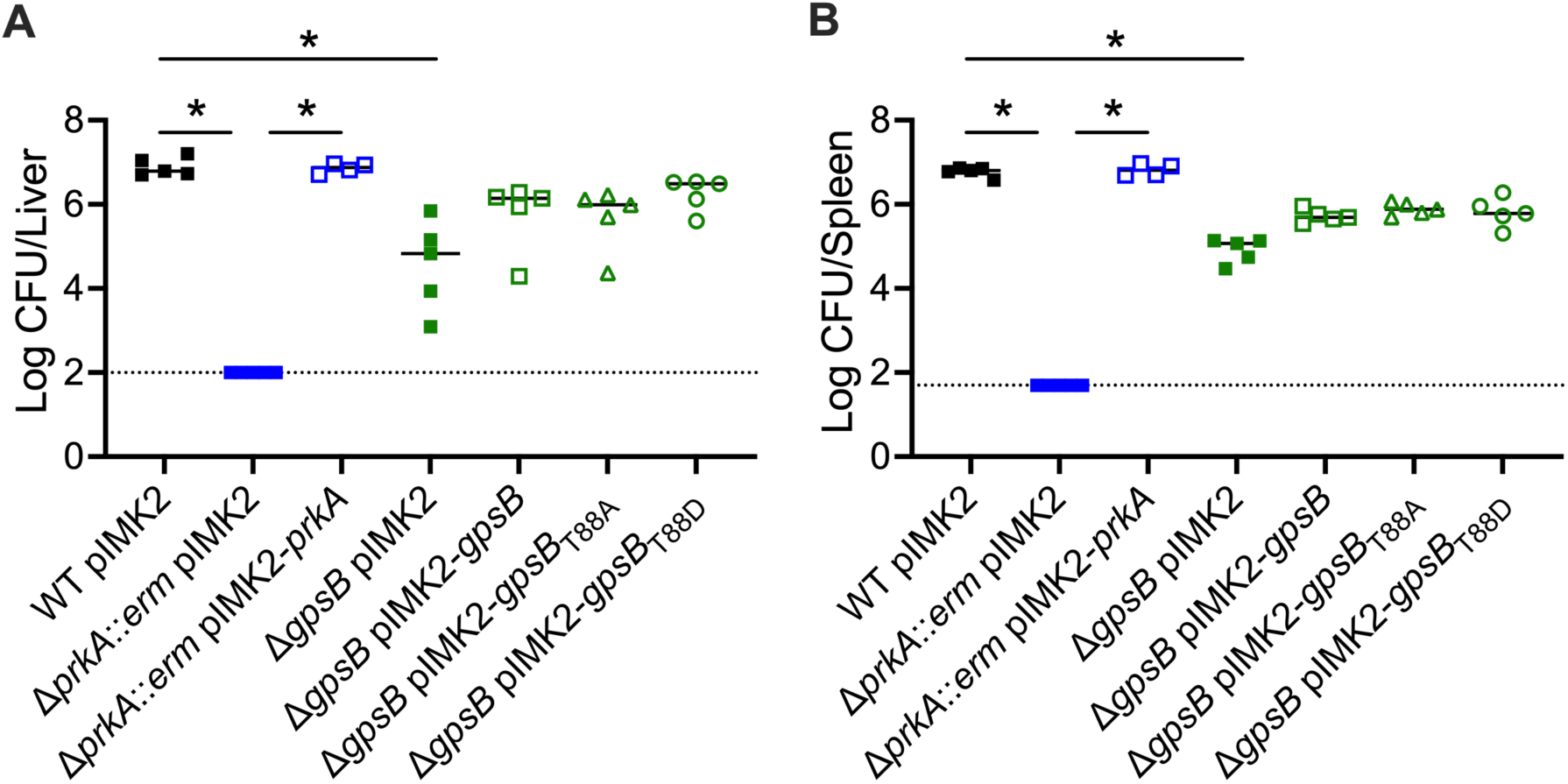
GpsB is required for systemic *L. monocytogenes* virulence due to PrkA-dependent and -independent activities. (**A**-**B**) Six-week-old C57BL/6 mice were infected intravenously with 1 x 10^5^ CFU of the indicated strains. Forty-eight hours post-infection, livers (**A**) and spleens (**B**) were harvested and CFU enumerated. Solid bars indicate medians and dotted lines indicate limits of detection. Data are representative of two independent experiments with 4-5 mice per group. *, *P* < 0.05 for the indicated comparisons by Kruskal-Wallis test with Dunn’s multiple comparisons test.

To determine whether the requirement of GpsB is due to its role in potentiating PrkA activity, we also assessed the virulence of the Δ*gpsB* mutant strain complemented with *gpsB*_T88A_ and *gpsB*_T88D_ alleles. Interestingly, bacterial burdens resulting from both strains were intermediate between wild-type and Δ*gpsB* mutant burdens, i.e., both alleles partially rescued loss of *gpsB* (Fig. 5). Cumulatively, these results suggests that GpsB is required for wild-type levels of virulence of *L. monocytogenes*, and that its importance is partially due to its role in potentiating PrkA activity.

## DISCUSSION

PASTA kinases are important for intrinsic cell wall stress responses and virulence in diverse bacteria, including in major pathogens like *L. monocytogenes*. However, the downstream mechanisms through which they influence physiology are less well understood. In this work, we investigated the importance in *L. monocytogenes* of PrkA-dependent phosphorylation of its substrate GpsB. We previously identified GpsB as a substrate of PrkA during cell wall stress (24), and herein we validate that GpsB is directly phosphorylated by PrkA *in vitro*. We further show that while a phosphoablative T88A allele of GpsB potentiates PrkA activity *in vitro* and facilitates PG synthesis, a T88D allele mimicking PrkA phosphorylation increases sensitivity to cell wall stress and inhibits a concomitant PrkA-dependent increase in PG synthesis in those conditions. These results indicate that similar to in *B. subtilis*, *S. pneumoniae*, and *E. faecalis*, GpsB contributes to modulating activity of the PASTA kinase in *L. monocytogenes*. In addition, these findings demonstrate another PrkA-dependent phosphoregulatory event in addition to regulation of PG synthesis through ReoM (24) is important for the response to cell wall stress and adaptation to the intracellular niche in this professional cytosolic pathogen.

Our findings support a model, first described in *B. subtilis* (32) and subsequently in *E. faecalis* (34), that GpsB and PrkA form a negative feedback loop. However, there are notable differences in this pathway in *L. monocytogenes* compared to others. In both *B. subtilis* and *E. faecalis*, the presence of GpsB is required for activation of the PASTA kinase, and in *E. faecalis* a Δ*gpsB* mutant is as sensitive to cephalosporin-induced cell wall stress as a PASTA kinase mutant (32, 34). In contrast, we find that GpsB activity is not required for PrkA activity *in vitro*, and a Δ*gpsB* mutant is less sensitive to cell wall stress than a Δ*prkA* mutant, suggesting that PrkA retains some activity in *L. monocytogenes* in the absence of GpsB. Another interesting difference is apparent between strains of *L. monocytogenes*: while in the EGD-e background a Δ*gpsB* mutant is unable to grow at 42°C (27), we find that in 10403s a Δ*gpsB* mutant has a significant defect but is not fully inhibited. The ability of GpsB_T88A_ to fully rescue growth of EGD-e at high temperature might suggest that the GpsB-PrkA interaction is more important for growth in this strain than in 10403s. This idea is consistent with prior differences noted in the essentiality of the ReoM-PrkA pathway, where PrkA-dependent phosphorylation of ReoM is essential in EGD-e (25) but not 10403s (24). Taken together, these findings suggest that GpsB-mediated control of PASTA kinases varies in different organisms and may vary even in different strains of the same species.

That the Δ*gpsB* mutant was more sensitive to cell wall-targeting agents (Table 1) but was able to synthesize wild-type levels of PG in the presence of stress (Fig. 3B) was unexpected. Given that GpsB has a role in directing the activity of PBP A1 in *L. monocytogenes* (27), one potential explanation for this apparent discrepancy could be that loss of GpsB prevents proper regulation and/or localization needed for optimal PBP A1 activity during ceftriaxone-induced stress. A previous study has found that PBP A1 localization is not different between wild type and a Δ*gpsB* mutant in *L. monocytogenes* (27); however, these studies were performed in the absence of cell wall stress, which may affect GpsB control of PBP A1. The phosphorylation status of GpsB might similarly affect its interaction with PBP A1. However, in *B. subtilis*, neither a phosphoablative nor phosphomimetic variant of GpsB affects the *in vitro* activity of its cognate PBP differently than wild-type GpsB (35). Yet, it remains a possibility that the phosphorylation state of *L. monocytogenes* GpsB may affect its interaction with PBP A1, and future studies are underway to determine how GpsB phosphorylation affects its interaction with this and other protein partners.

Another unexpected observation was that the Δ*gpsB* mutant complemented with *gpsB* or *gpsB*_T88A_ exhibited higher PG labeling, even in the absence of stress. We attempted to determine whether this increase in PG labeling was PrkA-dependent, but transduction of the Δ*prkA*::*erm* allele into strains harboring *trans*-encoded *gpsB*, *gpsB*_T88A_, and *gpsB*_T88D_ resulted frequently in suppressor formation at routine growth temperature, and therefore detailed analyses using these strains was not possible. In addition to the role of GpsB homologs in regulating PBP activity, a recent study by Hammond *et al.* (41) determined that GpsB also modulates wall teichoic acid synthesis in *B. subtilis*, and Schulz *et al.* recently demonstrated that changes in wall teichoic acid composition can affect resistance of *L. monocytogenes* to cell wall insults (42). Future studies are warranted to understand whether GpsB has PrkA-independent roles in resisting cell wall stress responses in *L. monocytogenes*.

In conclusion, this work demonstrates that the phosphorylation-based feedback loop between GpsB and PrkA in *L. monocytogenes* is important for control of PG synthesis, cytosolic survival, and virulence. These findings improve our understanding of how the listerial PASTA kinase is regulated and how such regulation may differ in diverse Gram-positive bacteria. Further investigation into serine/threonine kinase-mediated signaling networks will continue to advance our understanding of intrinsic resistance mechanisms to cell wall-targeting antibiotics and to bacterial pathogenesis.

## MATERIALS AND METHODS

### Ethics statement

All experiments involving animals were approved by the Institutional Animal Care and Use Committee of the University of Wisconsin-Madison.

### Bacterial strains, routine growth conditions, and cloning

*L*. *monocytogenes* was grown in brain heart infusion (BHI) broth or on BHI + 1.5% agar plates at 30°C unless otherwise indicated and was frozen in BHI + 40% glycerol. *E*. *coli* was routinely grown at 37°C in LB or on LB + 1.5% agar plates and frozen in LB + 40% glycerol. *E*. *coli* XL1- Blue was used for sub-cloning, SM10 or S17 were used for conjugating plasmids into *L*. *monocytogenes*, and Rosetta (DE3) pLysS was used for protein expression. Antibiotics were used at the following concentrations: 100 μg/ml carbenicillin, 10 μg/ml chloramphenicol, 200 μg/ml streptomycin, 30 μg/ml kanamycin, 50 μg/ml gentamicin.

The pGEX-6P-PrkA_1-338_ plasmid (Table 2) was constructed by a two-part ligation (NEB Quick Ligation kit) of BamHI- and NotI-linearized pGEX-6P and PCR-generated *prkA*_1-338_ amplified with primers AS21 and AS81 (Table 3) from 10403s genomic DNA. The pET28b-GpsB plasmid was constructed by a ligation of NdeI- and SalI-linearized pET28b and PCR-generated *gpsB* amplified with primers JLK131 and JLK132 from 10403s genomic DNA. The pET28b-GpsB_T88A_ plasmid was constructed by a ligation of NdeI- and SalI-linearized pET28b with a PCR- generated fragment amplified with primers JLK131 and JLK132 from a gBlock (IDT) of *gpsB* with a ACA261GCG mutation. Plasmid inserts were verified by Sanger sequencing.

**Table 2.** Plasmids used in this study.

| Name | Description | Citation |
| --- | --- | --- |
| pKSV7-*ΔgpsB* | Allelic exchange vector for making clean deletion of *gpsB* | This study |
| pIMK2-*prkA* | For expression of *prkA* from a constitutive promoter | (24) |
| pIMK2-*gpsB* | For expression of *gpsB* from a constitutive promoter | This work |
| pIMK2-*gpsB*<sub>T88A</sub> | For expression of *gpsB*<sub>T88A</sub> from a constitutive promoter | This study |
| pIMK2-*gpsB*<sub>T88D</sub> | For expression of *gpsB*<sub>T88D</sub> from a constitutive promoter | This study |
| pBHE573 | Luciferase reporter vector for bacteriolysis assays | (37) |
| pGEX-6P-PrkA<sub>1-338</sub> | For expression of the PrkA kinase domain (1-338) from *L. monocytogenes* | This study |
| pET28b-GpsB | For expression of GpsB from *L. monocytogenes* | This study |
| pET28b-GpsB<sub>T88A</sub> | For expression of GpsB<sub>T88A</sub> from *L. monocytogenes* | This study |

**Table 3.**
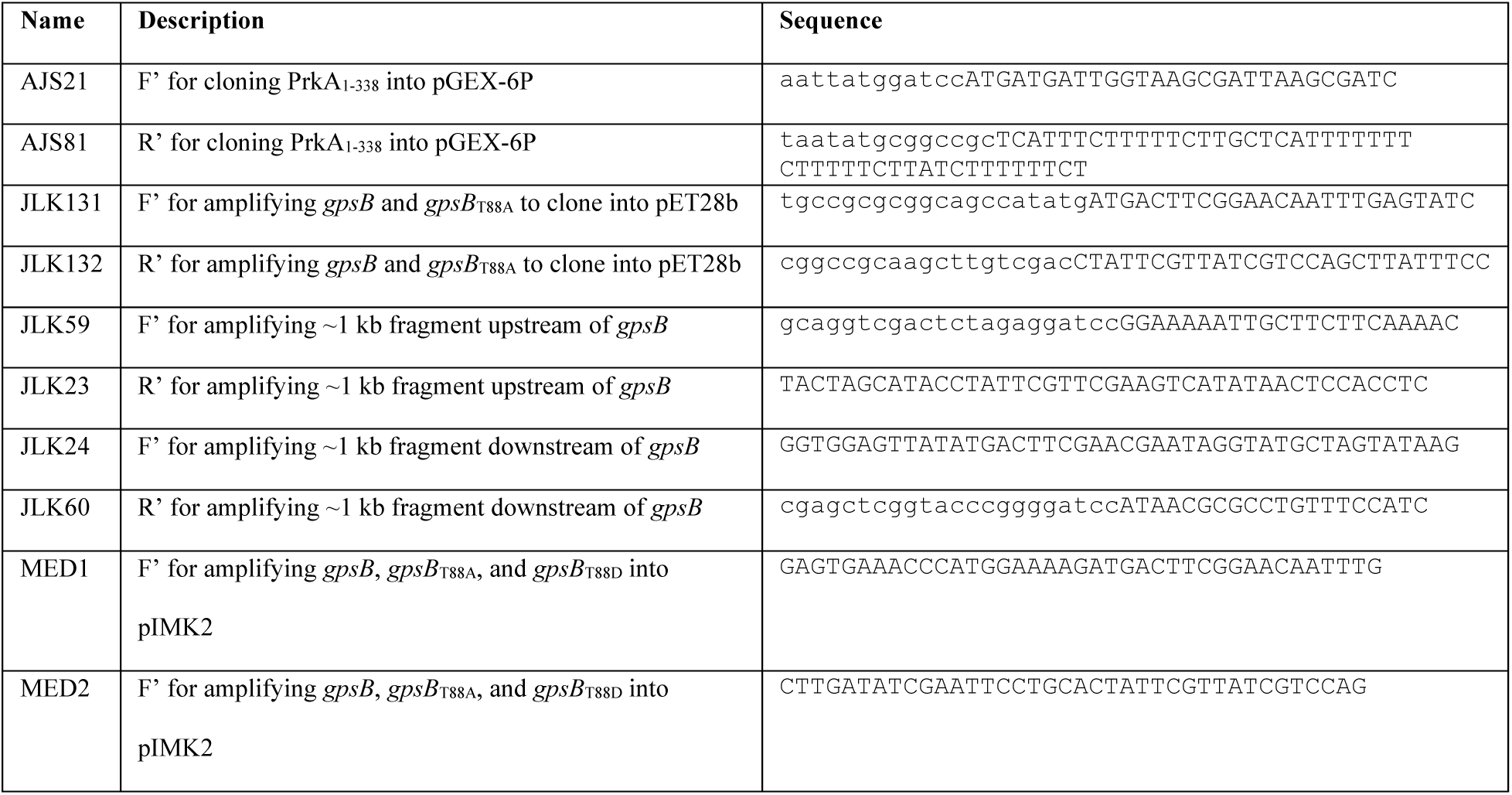
Primers used in this study.

The in-frame deletion of *gpsB* was generated in *L*. *monocytogenes* 10403s as previously described (43) using pKSV7-*oriT* (44). The pKSV7-Δ*gpsB* plasmid was constructed by Gibson Assembly with BamHI-linearized pKSV7-*oriT* and two PCR-generated fragments, a ∼1 kb fragment upstream of the *gpsB* ORF, amplified with primers JLK59 and JLK23, and a ∼1 kb fragment downstream of the *gpsB* ORF, amplified with primers JLK24 and JLK60 from *L*. *monocytogenes* 10403S gDNA. The pIMK2-*gpsB* plasmid was constructed by Gibson Assembly with BamHI- and PstI-linearized pIMK2 (45) and the *gpsB* ORF amplified with primers MED1 and MED2 from wild-type 10403s genomic DNA. The pIMK2-*gpsB*_T88A_ plasmid was constructed by Gibson Assembly with BamHI- and PstI-linearized pIMK2 and a PCR-generated fragment amplified with primers MED1 and MED2 from the *gpsB*_T88A_ gBlock described above. The pIMK2-*gpsB*_T88D_ plasmid was constructed by Gibson Assembly with BamHI- and PstI-linearized pIMK2 and a gBlock of the *gpsB* ORF with a ACA261GAG mutation and an additional 25 base pairs on the 5’ and 3’ end with homology to pIMK2. All plasmid inserts were verified by Sanger sequencing. For conjugations, donor *E. coli* SM10 or S17 and recipient *L. monocytogenes* strains were mixed at a 1:1 ratio on a non-selective BHI agar plate for 24 hours at 30°C. Transconjugants were selected by plating bacteria on BHI plates with streptomycin (for SM10 conjugations) or nalidixic acid (for S17 conjugations) plus chloramphenicol (pBHE573 and pKSV7-Δ*gpsB*) or kanamycin (pIMK2 and derivatives). Strains used in this study are listed in Table 4.

**Table 4.**
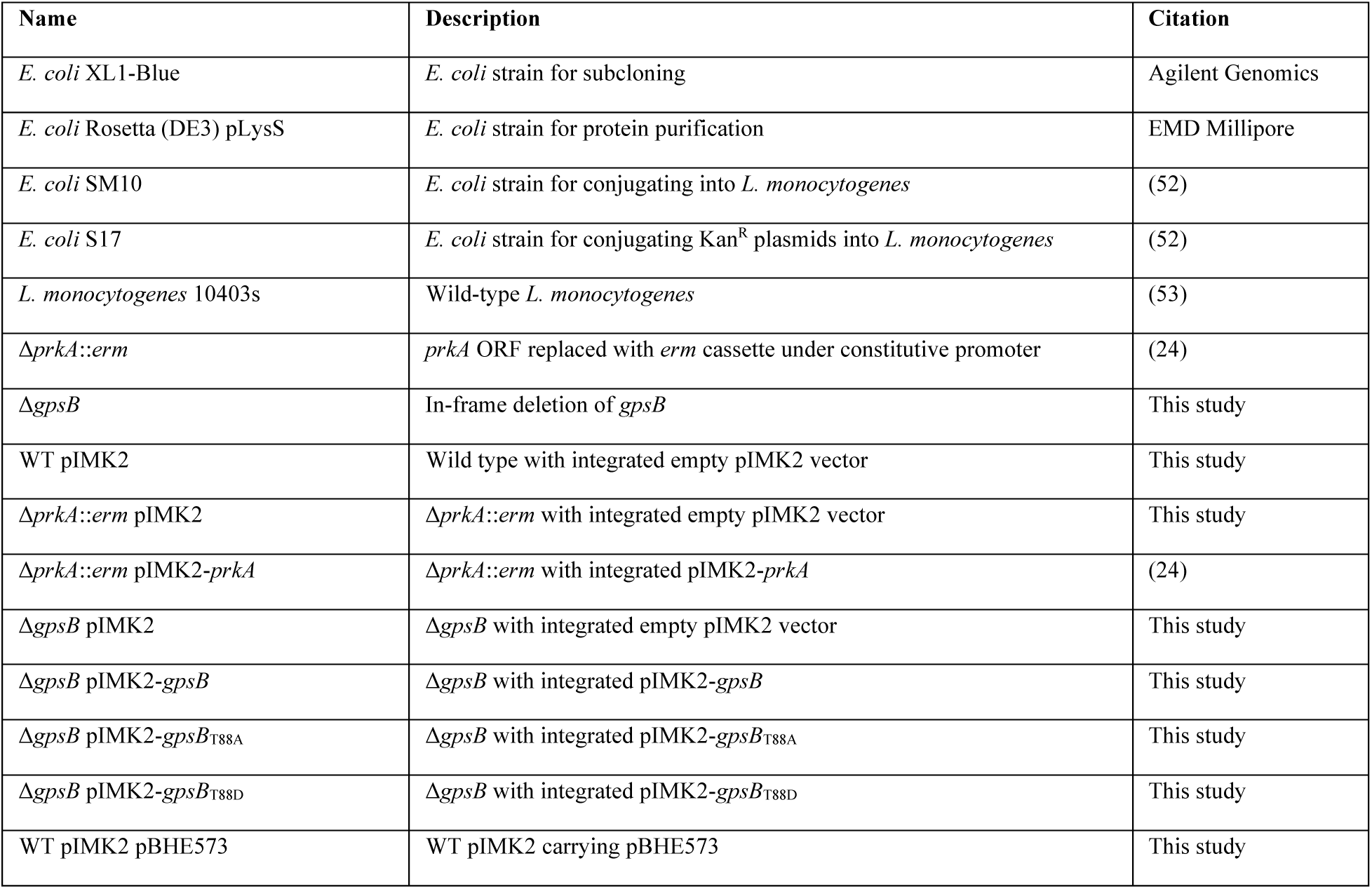
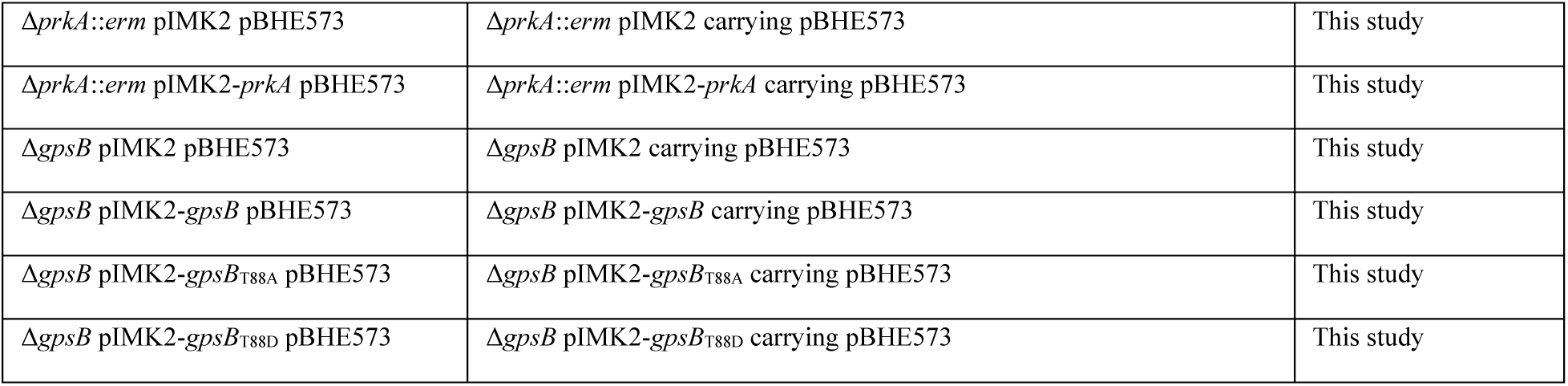
Strains used in this study.

### Protein purification

GST-tagged PrkA kinase domain plus linker region (PrkA_1-338_) was purified essentially as previously described (15, 16) with minor modifications: *E. coli* strain Rosetta (DE3) pLysS was used for protein production instead of BL21, a GSTrap column (Cytiva) was used instead of glutathione resin, and PreScission Protease was used to cleave the GST tag from PrkA_1-338._

His-tagged GpsB and GpsB_T88A_ were purified essentially as previously described (7) with minor modifications. Briefly, *E. coli* Rosetta (DE3) pLysS was freshly transformed with pET28b-GpsB or pET28b-GpsB_T88A_ and selected on LB agar plates with chloramphenicol and kanamycin. One transformant colony was used to inoculate 50 ml LB broth with chloramphenicol and kanamycin in a 250 ml baffled flask and grown at 37°C with shaking (250 RPM) overnight. The next morning, overnight cultures were diluted 1:100 into 250 ml LB broth with chloramphenicol and kanamycin in two separate 1 L baffled flasks and grown at 30°C with shaking (120 RPM) for 4 hours (OD_600_ ∼0.6). Isopropyl β-d-1-thiogalactopyranoside (IPTG) was added to a final concentration of 0.5 mM, and cultures were grown at 18°C with shaking (120 RPM) overnight (16 hours). The next morning, duplicate cultures were combined and centrifuged (3,220 x *g* at 4°C), and pellets were stored at -80°C. Pre- and post-induction samples were analyzed by SDS-PAGE. Cells were thawed on ice, resuspended in 20 ml PBS plus two cOmplete Mini EDTA-free protease inhibitor tablets (Roche) and 25 μg/ml phenylmethylsulfonyl fluoride (PMSF), and sonicated (12 cycles of 10 seconds on/10 seconds off) on ice. Lysates were clarified by centrifugation for 20 minutes at 16,000 x *g* at 4°C and subsequently filtered through a 0.22 μm filter. Half of the lysate (∼10 ml) was loaded onto a 1 ml HisTrap column (Cytiva) pre-equilibrated with PBS at 4°C, washed with 20 column volumes PBS, then washed with 20 column volumes PBS plus 25 mM imidazole, and then eluted in PBS plus 250 mM imidazole.

Eluted protein was dialyzed overnight in PBS at 4°C. Protein purity was assessed by SDS-PAGE and concentration was determined by BCA assay (Pierce) according to the manufacturer’s instructions. Protein aliquots were frozen in 10% glycerol at -80°C.

### *In vitro* kinase assays

Kinase assays were performed using the ATP analog ATPγS (46) with reaction conditions previously described (7). Endpoint kinase assays were performed by mixing 3 μg purified PrkA_1- 338_ with the indicated combinations of ∼3x molar excess of substrate to kinase (6 μg of the generic kinase substrate myelin basic protein [MBP; Novatein Biosciences, Woburn, MA]; 3 μg purified GpsB or GpsB_T88A_) in kinase buffer (50 mM Tris-HCl, pH 7.4; 0.5 mM dithiothreitol [DTT]; 5 mM MgCl_2_) plus 1 mM ATPγS (abcam, ab138911) in 30 μl total volume. Reactions were incubated overnight (∼16 hours) in the dark at room temperature. The next morning, 1.5 μl of 50 mM *p*-nitrobenzyl mesylate (PNBM; abcam, ab138910) in DMSO was added to each reaction and incubated for 2 hours in the dark at room temperature to alkylate thiophosphorylation sites. Samples were then subjected to Western blotting with an anti-thioester antibody (abcam92570) according to the manufacturer’s instructions. Briefly, SDS sample buffer was added to 1x, and samples were boiled at 95°C for 5 minutes. Samples were run on an SDS-PAGE gel, then transferred onto nitrocellulose using a semi-dry transfer apparatus. The membrane was blocked in TBS-T (TBS plus 0.5% Tween-20) containing 5% skim milk for 20 minutes, washed thrice for 5 minutes in TBS-T, and then incubated overnight (∼16 hours) in TBS-T with 5% skim milk plus a 1:5,000 dilution of anti-thioester antibody at 4°C, all with gentle rocking. The next morning, the membrane was washed thrice for 5 minutes in TBS-T, incubated for 1 hour at room temperature in TBS-T with 5% skim milk plus a 1:10,000 dilution of anti-rabbit DyLight 800 secondary (Rockland), and finally washed thrice for 5 minutes in TBS-T, all with gentle rocking. The membrane was immediately visualized using an Odyssey imager with ImageStudio software.

Kinetic kinase assays were performed by mixing 0.066 μM PrkA, 3.3 μM GpsB, and 14.2 μM MBP in kinase buffer plus 1 mM ATPγS in 60 μl total volume. ATPγS was added last, reactions were mixed by pipetting, and incubation was at room temperature in the dark. At the indicated time points, 15 μl aliquots were transferred to microcentrifuge tubes containing EDTA to quench the kinase reaction (final concentration of 10 mM EDTA), and 1.5 μl of 50 mM PNBM was added to each reaction and incubated for 2 hours in the dark at room temperature. Samples were then subjected to Western blotting with an anti-thioester antibody as detailed for endpoint assays above.

### Growth and minimum inhibitory concentration assays

*L. monocytogenes* strains were grown overnight (16–18 hours) in BHI at 30°C on a slant. For BHI growth assays, cultures were inoculated 1:50 into 100 μl of BHI in 96-well flat-bottom plates. For *L. monocytogenes* minimum inhibitory concentration (MIC) assays, cultures were inoculated 1:50 into 100 μl BHI containing serial 2-fold dilutions of the indicated compound in 96-well flat-bottom plates. Optical density (OD_600_) was measured every 15 minutes in an Eon Microplate Spectrophotometer (BioTek Instruments, Inc., Winooski, VT), and plates were incubated at the indicated temperature for growth assays or 37°C for MIC assays with 180 RPM double-orbital shaking between time points. The MIC was defined as the lowest concentration of a given compound at which the endpoint OD_600_ minus the starting OD_600_ was <0.1, the lowest OD_600_ with visible turbidity.

### Peptidoglycan labeling experiments

Peptidoglycan synthesis by *L. monocytogenes* grown in broth was measured by copper-catalyzed azide-alkyne cycloaddition (CuAAC) reactions as previously described (24, 36). Cells growing exponentially in BHI were back-diluted to OD_600_ 0.3 in 1 ml BHI medium, with or without the addition of 0.5 MIC of CRO (as determined in Fig. 3A). 0.5 mM of alkyne-D-alanine-D-alanine (EDA-DA, (47)) was added and cultures were grown for 1 hour to mid-log phase (final OD_600_ 0.6-1.0) at 37°C with shaking. 100 μl aliquots were washed twice with an equal volume PBS, fixed in ice-cold 70% ethanol for 10 minutes at -20°C, and again washed twice with PBS. Cells were resuspended in 50 μl of CuAAC reaction mixture (128 μM TBTA, 1 mM CuSO4, 1.2 mM freshly prepared sodium ascorbate, and 20 μM AFDye 488 Azide Plus fluorophore [Click Chemistry Tools] in PBS + 0.1% Triton X-100 + 0.01% BSA) and incubated with gentle rocking at room temperature for 30 minutes in the dark. Cells were washed twice with PBS and finally resuspended in FACS buffer. Samples were acquired using an Attune NxT flow cytometer with Attune Cytometric Software (Invitrogen) and analyzed using FlowJo software (Tree Star, Ashland, OR). Median fluorescence intensities were normalized to that of WT minus CRO.

### Macrophage cytosol killing assays

Intracellular lysis of *L. monocytogenes* was measured by luciferase reporter activity as previously described (37). Briefly, 5 x 10^5^ immortalized *Ifnar*^-/-^ BMDMs were seeded into 24- well plates overnight. *L. monocytogenes* strains carrying the pBHE573 reporter construct were grown overnight (16–18 hours) in BHI plus chloramphenicol at 30°C on a slant. Cells were infected at an MOI of 10. One hour post-infection, media was removed from cells and replaced with fresh media containing gentamicin. Six hours post-infection, cells were lysed, and luciferase activity was measured using luciferin reagent as previously described (37). Luminescence was measured in a Synergy HT Microplate Spectrophotometer (BioTek Instruments, Inc., Winooski, VT).

### Intramacrophage growth assays

Intracellular growth assays were performed as previously described (48) in bone marrow-derived macrophages (BMDMs) isolated as previously described (49). Briefly, 5 x 10^6^ cells were seeded in 60 mm dishes with coverslips overnight. *L. monocytogenes* strains were grown overnight (16–18 hours) in BHI at 30°C on a slant. BMDMs were infected at an MOI of 0.2. Thirty minutes post-infection, the media was removed, cells were washed twice with pre-warmed PBS, and then fresh media containing gentamicin was added. The number of CFUs per coverslip was determined at the indicated time points as previously described (50). Shown is a representative growth curve from three biological replicates.

### Mouse infection model

Intravenous mouse infections were performed as previously described (51). *L. monocytogenes* strains were grown overnight (16–18 hours) in BHI at 30°C on a slant. Overnight cultures were back-diluted 1:5 into BHI and grown at 37°C with shaking to mid-log phase (OD_600_ 0.4–0.6). Cultures were then washed with PBS and adjusted to 5 x 10^5^ CFU/ml. Six-week-old C57BL/6 mice (Charles River Laboratories) were infected with 1 x 10^5^ CFU in 200 μl PBS. Forty-eight hours post-infection, animals were euthanized by CO_2_ asphyxiation according to AVMA standards. Spleens and livers were harvested, homogenized in PBS + 0.1% NP-40, and plated for CFU. Infections were performed in biological duplicate, one with female mice and one with male mice.

## ACKNOWLEDGMENTS

The authors thank Dr. Adam Schaenzer for constructing the pGEX-6P-PrkA1-338 plasmid.

This work was supported by National Institutes of Health grants awarded to J.L.K. (F32- AI154570) and J.-D.S. (R01-AI137070 and R21-AI144060) and a Burroughs Wellcome Fund Investigators in the Pathogenesis of Infectious Diseases Award awarded to J.-D.S. The funders had no role in study design, data collection and analysis, decision to publish, or preparation of the manuscript.

## AUTHOR CONTRIBUTIONS

J.L.K. and M.E.D. performed the research. J.L.K. and J.-D.S. designed the experiments, analyzed the data, and wrote the paper. All authors approved the final version for submission.

## REFERENCES

1. World Health Organization. 2022. Global antimicrobial resistance and use surveillance system (GLASS) report 2022. Geneva.

2. Pensinger DA, Schaenzer AJ, Sauer JD. 2018. Do Shoot the Messenger: PASTA Kinases as Virulence Determinants and Antibiotic Targets. Trends Microbiol 26:56–69.

3. Ohlsen K, Donat S. 2010. The impact of serine/threonine phosphorylation in *Staphylococcus aureus*. International Journal of Medical Microbiology 300:137–141.

4. Manuse S, Fleurie A, Zucchini L, Lesterlin C, Grangeasse C. 2016. Role of eukaryotic-like serine/threonine kinases in bacterial cell division and morphogenesis. FEMS Microbiol Rev 40.

5. Prust N, van der Laarse S, van den Toorn HWP, van Sorge NM, Lemeer S. 2021. In-Depth Characterization of the *Staphylococcus aureus* Phosphoproteome Reveals New Targets of Stk1. Molecular & Cellular Proteomics 20.

6. Iannetta AA, Minton NE, Uitenbroek AA, Little JL, Stanton CR, Kristich CJ, Hicks LM. 2021. IreK-Mediated, Cell Wall-Protective Phosphorylation in *Enterococcus faecalis*. J Proteome Res 20:5131–5144.

7. Pensinger DA, Boldon KM, Chen GY, Vincent WJ, Sherman K, Xiong M, Schaenzer AJ, Forster ER, Coers J, Striker R, Sauer JD. 2016. The *Listeria monocytogenes* PASTA Kinase PrkA and Its Substrate YvcK Are Required for Cell Wall Homeostasis, Metabolism, and Virulence. PLoS Pathog 12:e1006001.

8. Debarbouille M, Dramsi S, Dussurget O, Nahori MA, Vaganay E, Jouvion G, Cozzone A, Msadek T, Duclos B. 2009. Characterization of a serine/threonine kinase involved in virulence of Staphylococcus aureus. J Bacteriol 191:4070–4081.

9. Tamber S, Schwartzman J, Cheung AL. 2010. Role of PknB Kinase in Antibiotic Resistance and Virulence in Community-Acquired Methicillin-Resistant *Staphylococcus aureus* Strain USA300. Infect Immun 78.

10. Kristich CJ, Little JL, Hall CL, Hoff JS. 2011. Reciprocal regulation of cephalosporin resistance in *Enterococcus faecalis*. mBio 2:e00199–11.

11. Echenique J, Kadioglu A, Romao S, Andrew PW, Trombe MC. 2004. Protein serine/threonine kinase StkP positively controls virulence and competence in Streptococcus pneumoniae. Infect Immun 72:2434–2437.

12. Dias R, Félix D, Caniça M, Trombe M-C. 2009. The highly conserved serine threonine kinase StkP of *Streptococcus pneumoniae* contributes to penicillin susceptibility independently from genes encoding penicillin-binding proteins. BMC Microbiol 9.

13. Chawla Y, Upadhyay S, Khan S, Nagarajan SN, Forti F, Nandicoori VK. 2014. Protein kinase B (PknB) of Mycobacterium tuberculosis is essential for growth of the pathogen in vitro as well as for survival within the host. J Biol Chem 289:13858–13875.

14. Bugrysheva J, Froehlich BJ, Freiberg JA, Scott JR. 2011. Serine/Threonine Protein Kinase Stk Is Required for Virulence, Stress Response, and Penicillin Tolerance in *Streptococcus pyogenes*. Infect Immun 79.

15. Pensinger DA, Aliota MT, Schaenzer AJ, Boldon KM, Ansari IU, Vincent WJ, Knight B, Reniere ML, Striker R, Sauer JD. 2014. Selective pharmacologic inhibition of a PASTA kinase increases *Listeria monocytogenes* susceptibility to beta-lactam antibiotics. Antimicrob Agents Chemother 58:4486–4494.

16. Schaenzer AJ, Wlodarchak N, Drewry DH, Zuercher WJ, Rose WE, Striker R, Sauer JD. 2017. A screen for kinase inhibitors identifies antimicrobial imidazopyridine aminofurazans as specific inhibitors of the *Listeria monocytogenes* PASTA kinase PrkA. J Biol Chem 292:17037–17045.

17. Schaenzer AJ, Wlodarchak N, Drewry DH, Zuercher WJ, Rose WE, Ferrer CA, Sauer JD, Striker R. 2018. GW779439X and Its Pyrazolopyridazine Derivatives Inhibit the Serine/Threonine Kinase Stk1 and Act As Antibiotic Adjuvants against beta-Lactam-Resistant *Staphylococcus aureus*. ACS Infect Dis 4:1508–1518.

18. Kant S, Asthana S, Missiakas D, Pancholi V. 2017. A novel STK1-targeted small-molecule as an “antibiotic resistance breaker” against multidrug-resistant *Staphylococcus aureus*. Sci Rep 7.

19. de Noordhout CM, Devleesschauwer B, Angulo FJ, Verbeke G, Haagsma J, Kirk M, Havelaar A, Speybroeck N. 2014. The global burden of listeriosis: a systematic review and meta-analysis. Lancet Infect Dis 14:1073–1082.

20. Scallan E, Hoekstra RM, Angulo FJ, Tauxe R V, Widdowson MA, Roy SL, Jones JL, Griffin PM. 2011. Foodborne illness acquired in the United States--major pathogens. Emerg Infect Dis 17:7–15.

21. Radoshevich L, Cossart P. 2018. *Listeria monocytogenes*: towards a complete picture of its physiology and pathogenesis. Nat Rev Microbiol 16:32–46.

22. Goetz M, Bubert A, Wang G, Chico-Calero I, Vazquez-Boland JA, Beck M, Slaghuis J, Szalay AA, Goebel W. 2001. Microinjection and growth of bacteria in the cytosol of mammalian host cells. Proc Natl Acad Sci U S A 98:12221–12226.

23. Randow F, MacMicking JD, James LC. 2013. Cellular Self-Defense: How Cell-Autonomous Immunity Protects Against Pathogens. Science (1979) 340.

24. Kelliher JL, Grunenwald CM, Abrahams RR, Daanen ME, Lew CI, Rose WE, Sauer J-D. 2021. PASTA kinase-dependent control of peptidoglycan synthesis via ReoM is required for cell wall stress responses, cytosolic survival, and virulence in *Listeria monocytogenes*. PLoS Pathog 17:e1009881.

25. Wamp S, Rutter ZJ, Rismondo J, Jennings CE, Möller L, Lewis RJ, Halbedel S. 2020. PrkA controls peptidoglycan biosynthesis through the essential phosphorylation of ReoM. Elife 9.

26. Claessen D, Emmins R, Hamoen LW, Daniel RA, Errington J, Edwards DH. 2008. Control of the cell elongation–division cycle by shuttling of PBP1 protein in *Bacillus subtilis*. Mol Microbiol 68:1029–1046.

27. Rismondo J, Cleverley RM, Lane H V., Großhennig S, Steglich A, Möller L, Mannala GK, Hain T, Lewis RJ, Halbedel S. 2016. Structure of the bacterial cell division determinant GpsB and its interaction with penicillin-binding proteins. Mol Microbiol 99:978–998.

28. Cleverley RM, Rutter ZJ, Rismondo J, Corona F, Tsui H-CT, Alatawi FA, Daniel RA, Halbedel S, Massidda O, Winkler ME, Lewis RJ. 2019. The cell cycle regulator GpsB functions as cytosolic adaptor for multiple cell wall enzymes. Nat Commun 10:261.

29. Fleurie A, Manuse S, Zhao C, Campo N, Cluzel C, Lavergne J-P, Freton C, Combet C, Guiral S, Soufi B, Macek B, Kuru E, VanNieuwenhze MS, Brun Y V., Di Guilmi A-M, Claverys J-P, Galinier A, Grangeasse C. 2014. Interplay of the Serine/Threonine-Kinase StkP and the Paralogs DivIVA and GpsB in Pneumococcal Cell Elongation and Division. PLoS Genet 10:e1004275.

30. Rued BE, Zheng JJ, Mura A, Tsui H-CT, Boersma MJ, Mazny JL, Corona F, Perez AJ, Fadda D, Doubravová L, Buriánková K, Branny P, Massidda O, Winkler ME. 2017. Suppression and synthetic-lethal genetic relationships of Δ*gpsB* mutations indicate that GpsB mediates protein phosphorylation and penicillin-binding protein interactions in *Streptococcus pneumoniae* D39. Mol Microbiol 103:931–957.

31. Rismondo J, Bender JK, Halbedel S. 2017. Suppressor Mutations Linking *gpsB* with the First Committed Step of Peptidoglycan Biosynthesis in *Listeria monocytogenes*. J Bacteriol 199.

32. Pompeo F, Foulquier E, Serrano B, Grangeasse C, Galinier A. 2015. Phosphorylation of the cell division protein GpsB regulates PrkC kinase activity through a negative feedback loop in *Bacillus subtilis*. Mol Microbiol 97.

33. Fleurie A, Manuse S, Zhao C, Campo N, Cluzel C, Lavergne J-P, Freton C, Combet C, Guiral S, Soufi B, Macek B, Kuru E, VanNieuwenhze MS, Brun Y v., di Guilmi A-M, Claverys J-P, Galinier A, Grangeasse C. 2014. Interplay of the Serine/Threonine-Kinase StkP and the Paralogs DivIVA and GpsB in Pneumococcal Cell Elongation and Division. PLoS Genet 10:e1004275.

34. Minton NE, Djorić D, Little J, Kristich CJ. 2022. GpsB Promotes PASTA Kinase Signaling and Cephalosporin Resistance in *Enterococcus faecalis*. J Bacteriol 204.

35. Cleverley RM, Rismondo J, Lockhart-Cairns MP, Van Bentum PT, Egan AJF, Vollmer W, Halbedel S, Baldock C, Breukink E, Lewis RJ. 2016. Subunit Arrangement in GpsB, a Regulator of Cell Wall Biosynthesis. Microbial Drug Resistance 22:446–460.

36. Siegrist MS, Whiteside S, Jewett JC, Aditham A, Cava F, Bertozzi CR. 2013. d-Amino Acid Chemical Reporters Reveal Peptidoglycan Dynamics of an Intracellular Pathogen. ACS Chem Biol 8.

37. Sauer JD, Witte CE, Zemansky J, Hanson B, Lauer P, Portnoy DA. 2010. *Listeria monocytogenes* triggers AIM2-mediated pyroptosis upon infrequent bacteriolysis in the macrophage cytosol. Cell Host Microbe 7:412–419.

38. Chen GY, McDougal CE, D’Antonio MA, Portman JL, Sauer JD. 2017. A Genetic Screen Reveals that Synthesis of 1,4-Dihydroxy-2-Naphthoate (DHNA), but Not Full-Length Menaquinone, Is Required for *Listeria monocytogenes* Cytosolic Survival. mBio 8.

39. Beuzón CR, Salcedo SP, Holden DW. 2002. Growth and killing of a *Salmonella enterica* serovar Typhimurium *sifA* mutant strain in the cytosol of different host cell lines. Microbiology (N Y) 148.

40. Laguna RK, Creasey EA, Li Z, Valtz N, Isberg RR. 2006. A *Legionella pneumophila*- translocated substrate that is required for growth within macrophages and protection from host cell death. Proceedings of the National Academy of Sciences 103.

41. Hammond LR, Sacco MD, Khan SJ, Spanoudis C, Hough-Neidig A, Chen Y, Eswara PJ. 2022. GpsB Coordinates Cell Division and Cell Surface Decoration by Wall Teichoic Acids in *Staphylococcus aureus*. Microbiol Spectr 10.

42. Schulz LM, Rothe P, Halbedel S, Gründling A, Rismondo J. 2022. Imbalance of peptidoglycan biosynthesis alters the cell surface charge of *Listeria monocytogenes*. The Cell Surface 8:100085.

43. Camilli A, Tilney LG, Portnoy DA. 1993. Dual roles of *plcA* in *Listeria monocytogenes* pathogenesis. Mol Microbiol 8.

44. Smith K, Youngman P. 1992. Use of a new integrational vector to investigate compartment-specific expression of the *Bacillus subtilis spollM* gene. Biochimie 74.

45. Monk IR, Gahan CGM, Hill C. 2008. Tools for Functional Postgenomic Analysis of *Listeria monocytogenes*. Appl Environ Microbiol 74.

46. Allen JJ, Li M, Brinkworth CS, Paulson JL, Wang D, Hübner A, Chou W-H, Davis RJ, Burlingame AL, Messing RO, Katayama CD, Hedrick SM, Shokat KM. 2007. A semisynthetic epitope for kinase substrates. Nat Methods 4:511–516.

47. Liechti GW, Kuru E, Hall E, Kalinda A, Brun Y v., VanNieuwenhze M, Maurelli AT. 2014. A new metabolic cell-wall labelling method reveals peptidoglycan in *Chlamydia trachomatis*. Nature 506.

48. Smith HB, Li TL, Liao MK, Chen GY, Guo Z, Sauer J-D. 2021. *Listeria monocytogenes* MenI encodes a DHNA-CoA thioesterase necessary for menaquinone biosynthesis, cytosolic survival, and virulence. Infect Immun https://doi.org/10.1128/IAI.00792-20.

49. Jones S, Portnoy DA. 1994. Characterization of *Listeria monocytogenes* pathogenesis in a strain expressing perfringolysin O in place of listeriolysin O. Infect Immun 62.

50. Portnoy DA, Jacks PS, Hinrichs DJ. 1988. Role of hemolysin for the intracellular growth of *Listeria monocytogenes*. Journal of Experimental Medicine 167.

51. Sauer JD, Pereyre S, Archer KA, Burke TP, Hanson B, Lauer P, Portnoy DA. 2011. *Listeria monocytogenes* engineered to activate the Nlrc4 inflammasome are severely attenuated and are poor inducers of protective immunity. Proc Natl Acad Sci U S A 108:12419–12424.

52. Simon R, Priefer U, Pühler A. 1983. A Broad Host Range Mobilization System for In Vivo Genetic Engineering: Transposon Mutagenesis in Gram Negative Bacteria. Bio/Technology 1:784–791.

53. Bécavin C, Bouchier C, Lechat P, Archambaud C, Creno S, Gouin E, Wu Z, Kühbacher A, Brisse S, Pucciarelli MG, García-del Portillo F, Hain T, Portnoy DA, Chakraborty T, Lecuit M, Pizarro-Cerdá J, Moszer I, Bierne H, Cossart P. 2014. Comparison of Widely Used *Listeria monocytogenes* Strains EGD, 10403S, and EGD-e Highlights Genomic Differences Underlying Variations in Pathogenicity. mBio 5.

